# Self-Regulation of Visual Word Form Area activation with real-time fMRI neurofeedback

**DOI:** 10.1101/2022.11.25.517926

**Authors:** Amelie Haugg, Nada Frei, Milena Menghini, Felizia Stutz, Sara Steinegger, Martina Röthlisberger, Silvia Brem

## Abstract

The Visual Word Form Area (VWFA) is a key region of the brain’s reading network and its activation has been shown to be strongly associated with reading skills. Here, for the first time, we investigated whether voluntary regulation of VWFA activity is feasible using real-time fMRI neurofeedback. 40 adults with typical reading skills were instructed to either upregulate (UP group) or downregulate (DOWN group) their own VWFA activity during six neurofeedback training runs. The VWFA target region was individually defined based on a functional localizer task. Before and after training, also regulation runs without feedback (“no-feedback runs”) were performed.

When comparing the two groups, we found stronger activity across the whole reading network, including the VWFA, for the UP than the DOWN group. Crucially, we observed a significant interaction of group and time (pre, post) for the no-feedback runs: The two groups did not differ in their VWFA activity before neurofeedback training, but the UP group showed significantly stronger activity than the DOWN group after neurofeedback training. Our results indicate that self-regulation of the VWFA activity is feasible and that, once learned, successful self-regulation can even be performed in the absence of feedback. These results are a crucial step toward to development of a potential clinical intervention to improve reading skills in individuals with reading impairments.

## Introduction

Reading is a key skill in our society and substantially influences academic and socio-economic development (Slavin et al., 2009). As a consequence, individuals with poor reading skills, such as individuals with developmental dyslexia, suffer from disadvantages in their education and employment (Frederickson & Jacobs, 2001; Tanner, 2009), and demonstrate lower scores in overall quality of life (Y. Huang et al., 2020; Kalka & Lockiewicz, 2018). In the brain, reading skills have been associated with the functionality of a diverse set of brain regions such as the inferior frontal gyrus (IFG), the intraparietal lobule (IPL), the superior temporal gyrus (STG), the left precentral gyrus (PCG), and the ventral occipitotemporal cortex (vOTC) (Cao, 2016; Romanovska & Bonte, 2021) which together form the brain’s reading network (Martin et al., 2015).

One key structure of the reading network responsible for the processing of written stimuli is the Visual Word Form System along the fusiform gyrus in the ventral occipito-temporal cortex, with the Visual Word Form Area (VWFA) located in the left mid-fusiform gyrus as its core. The VWFA is suggested to process print on hierarchical posterior-to-anterior and lateral-to-medial axes, with more recent studies suggesting a subdivision of VWFA functions (Bouhali et al., 2019; Caffarra et al., 2021; Lerma-Usabiaga et al., 2018a; Vinckier et al., 2007). More posterior regions of the VWFA thus receive bottom-up information from the visual cortex and are particularly responsive to the presentation of orthographic stimuli and perceptual aspects of print. Anterior portions of the VWFA integrate top-down information from higher-order areas of the reading and language network and are related to lexical processing (Caffarra et al., 2021; Lerma-Usabiaga et al., 2018).

Importantly, VWFA activity levels have been shown to be associated with reading skills (Brem et al., 2020; Dehaene et al., 2010). Both poor-reading and illiterate individuals were found to show lower VWFA activity than typical readers (Maisog et al., 2008; Richlan et al., 2009). Furthermore, also lesion studies in adults support the strong interrelation between reading performance and VWFA function (Pflugshaupt et al., 2009; P. E. Turkeltaub et al., 2014). This finding makes the VWFA a promising target for interventions aiming at improving orthographic processing. Importantly, Hirshorn and colleagues also demonstrated causality between VWFA activation and reading performance (Hirshorn et al., 2016). They showed that disruption of VWFA activity with intracranial electrodes led to an impaired perception of words and letters. In accordance, a normalization of VWFA signals through brain-based interventions might also support reading.

One method which allows for a fast and long-lasting regulation of brain signals in order to normalize dysfunctional neural signals is real-time fMRI neurofeedback (rtfMRI NF). rtfMRI NF enables its user to voluntarily control their own brain signals within or between predefined regions of interest (ROIs) by providing feedback on ongoing brain measures acquired with an MRI scanner (Sitaram et al., 2016; Sulzer, Haller, et al., 2013; Weiskopf, 2012). Using this feedback information, the user can adapt their own mental processes (e.g. mental strategies) to direct the acquired brain signals in the desired direction. In the past, rtfMRI NF has been applied to a wide range of different target ROIs, including subcortical regions such as the amygdala (Brühl et al., 2014; Hellrung et al., 2018; Young et al., 2014) or the ventral tegmental area (Kirschner et al., 2018; MacInnes et al., 2016; Sulzer, Sitaram, et al., 2013), and cortical regions such as the auditory cortex (Emmert et al., 2017), supplementary motor area (Liew et al., 2016), or the visual cortex (Scharnowski et al., 2012). Importantly, Pereira and colleagues already demonstrated that successful self-regulation of activity within the bilateral fusiform face area, which lies in a close location to the VWFA in the left hemisphere, is indeed feasible (Pereira et al., 2019). Further, rtfMRI NF has been demonstrated to successfully improve behavioral measures linked to the targeted ROIs, such as attention (Pamplona et al., 2020), memory (Scharnowski et al., 2015), motivation (Zhi et al., 2018), and visual perception (Scharnowski et al., 2012). Importantly, the normalization of dysfunctional measures has also been shown to lead to significant improvements in clinical measures for patients suffering from disorders such as depression (Linden et al., 2012; Young et al., 2017), post-traumatic stress disorder (Nicholson et al., 2017), or Huntington’s disease (Papoutsi et al., 2019).

To date, however, rtfMRI NF has never been used in the context of reading and it has not been used to target the VWFA activation. NF training of the VWFA might constitute a promising intervention for individuals with reading impairments to normalize their hypoactive VWFA activity with the ultimate goal of improving reading skills. Here, we conducted a first study to investigate the feasibility of up- and downregulation of the VWFA in healthy individuals. In addition, we investigated whether, once learned, self-regulation of the VWFA would also be possible to perform in the absence of feedback.

## Methods

### Participants

45 right-handed adults were recruited for the study. They were randomly assigned to either an upregulation (UP) or downregulation (DOWN) group. Inclusion criteria included typical reading performance (reading fluency scores higher than the 30^th^ percentile for both words and pseudowords) and non-verbal IQ scores above a threshold of 85, which resulted in the exclusion of four participants due to low performance in reading fluency. Further exclusion criteria were psychiatric or neurological disorders according to DSM-5 (American Psychiatric Association, 2013) and MRI-incompatibility (e.g. metal implants, pacemakers, claustrophobia, pregnancy). One additional participant was excluded due to an incidental finding, leaving a total of 20 participants in the UP group and 20 participants in the DOWN group. A detailed description of demographic, cognitive, and reading scores for the UP and DOWN group is provided in Table 1 in the results section. All participants provided written informed consent and were compensated 60 CHF for their participation. The study was approved by the ethics committee of the Kanton of Zurich.

**Table 1:** Overview of demographic and behavioral measures in the UP and DOWN participant groups.

|  | **UP group:**<br>**mean, std** | **DOWN group:**<br>**mean, std** | **Group difference** |
| --- | --- | --- | --- |
| **Age** | 26.76, 5.144 | 25.54, 3.60 | t(38)=0.87, p=0.39 |
| **Sex** | 8m, 12f | 10m, 10f |  |
| **School years** | 11.63, 1.3 | 12.2, 1.47 | t(37)=1.28, p=0.21 |
| **Non-verbal**<br>**intelligence (RIAS)** | 109.75, 5.58 | 110.68, 3.72 | t(33)=0.59, p=0.56 |
| **Working memory**<br>**(WAIS)** | 20.30, 4.21 | 19.00, 2.70 | t(38)=1.16, p=0.25 |
| **Handedness (EHI)** | 78.18, 15.51 | 83.33, 17.56 | t(37)=0.34, p=0.34 |
| **Mental imagery**<br>**(VVIQ)** | 35.42, 10.17 | 37.5, 13.61 | t(37)=0.54, p=0.59 |
| **Motivation** | 8.65, 1.23 | 7.95, 1.85 | t(38)=1.41, p=0.17 |
| **Tiredness before**<br>**training** | 3.70, 1.84 | 3.00, 1.03 | t(38)=1.49, p=0.15 |
| **Reading**<br>**comprehension**<br>**(LGVT)** | 57.45, 24.44 | 61.55, 30.10 | t(38)=0.47, p=0.64 |
| **Reading speed**<br>**(LGVT)** | 49.65, 25.41 | 55.10, 31.38 | t(38)=0.60, p=0.55 |
| **Reading accuracy**<br>**(LGVT)** | 71.35, 23.91 | 74.45, 26.97 | t(38)=0.39, p=0.70 |
| **Reading fluency**<br>**words (SLRT-II)** | 64.16, 18.39 | 72.33, 12.44 | t(34)=1.59, p=0.12 |
| **Reading fluency**<br>**pseudowords**<br>**(SLRT-II)** | 69.81, 16.32 | 75.85, 15.91 | t(34)=1.12, p=0.27 |
| **Reading fluency**<br>**overall (SLRT-II)** | 66.98, 14.78 | 74.09, 11.53 | t(34)=1.62, p=0.11 |

### Screening and behavioral testing

Prior to study inclusion, all volunteers were screened via telephone. Participants who fulfilled the basic inclusion criteria were then provided with a set of questionnaires which they filled out online at home using RedCap (https://www.project-redcap.org/). These questionnaires included the Edinburgh Handedness Inventory (EHI) to assess the level of handedness (Oldfield, 1971), the Adult Reading History Questionnaire (Lefly & Pennington, 2000), the Vividness of Visual Imagery Questionnaire to get information on each participant’s abilities to perform vivid mental imagery (VVIQ) (Marks, 1973), and a custom questionnaire on language skills and basic demographic information.

In addition, participants performed several cognitive tests online during a 20 min video call session with an investigator. These tests assessed working memory using the forward and backward digit span test from the Wechsler Adult Intelligence Scale (WAIS-IV) (Drozdick et al., 2018; Wechsler, 1955), and non-verbal intelligence using two subtests of the German version of the Reynolds Intellectual Assessment Scale (RIAS, subtests *Odd-Item Out* and *What’s missing?)* (Reynolds & Kamphaus, 2003). Five participants reported being already familiar with the RIAS and were excluded from analyses concerning non-verbal intelligence.

Finally, participants underwent two reading tests in person at the MR Center of the Psychiatric University Hospital Zurich. Here, participants first performed a standardized reading fluency test (SLRT-II) (Moll & Landerl, 2010) where words and pseudowords had to be read out loud as fast and as accurately as possible within one minute. Then, participants underwent a standardized silent reading assessment of a longer text (LGVT) (Schneider et al., 2007) to measure reading comprehension, reading accuracy, and reading speed. In specific, participants were instructed to read as much of the text as possible within a time window of 6 minutes and to pick the correct answer from single-choice word options embedded in the text. The two reading tests took approximately 10 minutes and were repeated a second time after the MR session. Two versions of LGVT, which were randomly assigned to the participants, were used before and after the MR session to control for memory effects. Four participants reported being already familiar with the SLRT-II and were excluded from analyses involving reading fluency scores.

### Setup of the MR session

After behavioral testing, participants received instructions on the MR session. The MR session lasted approximately 1.5 hours, including a break outside the MR scanner after around one hour. The session included a functional VWFA localizer and a no-feedback run each before and after rtFMRI NF training as well as six rtfMRI NF training runs. An overview of the whole experimental design including the MR session is given in Figure 1.

**Figure 1:**
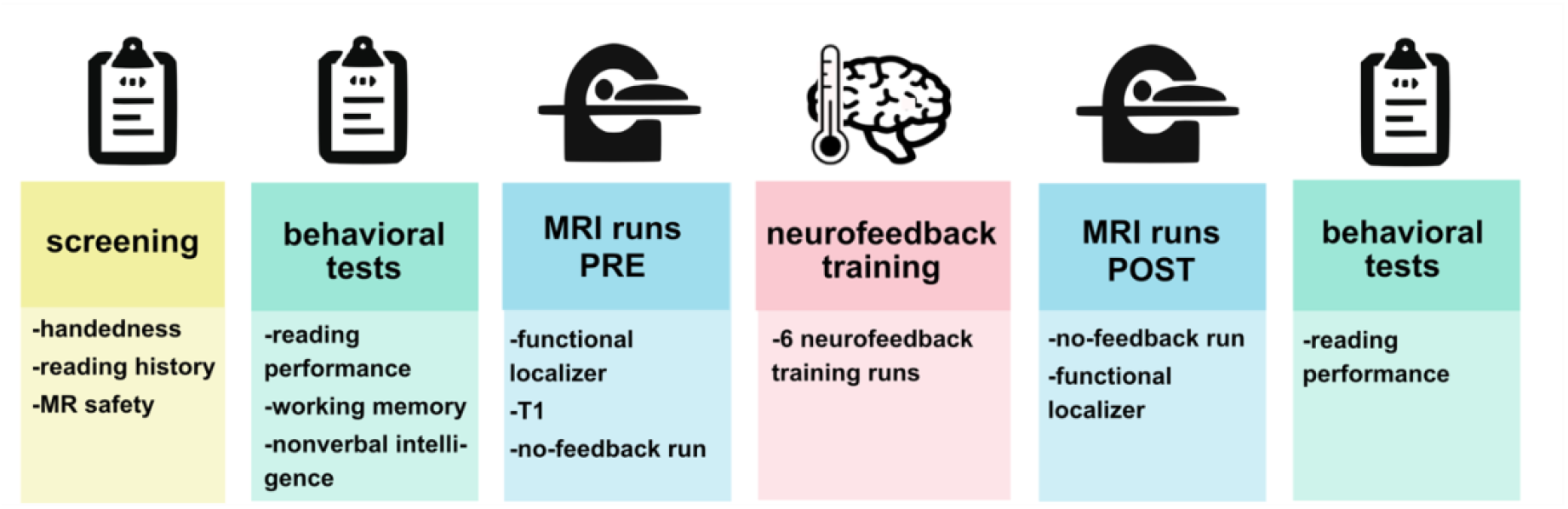
Experimental design. After an initial screening, participants underwent a set of behavioral tests on cognition and reading skills. During the MR session, participants first underwent a functional VWFA localizer, a T1-weighted anatomical scan, and a no-feedback run. Then, 6 runs of neurofeedback training were performed, followed by a break outside the scanner. After the break, participants repeated the no-feedback and functional localizer run in the MR scanner and the reading performance tests outside the scanner.

### Functional VWFA localizer runs

At the beginning of each MR session, a functional localizer scan was performed to identify the individual location of the VWFA for each participant and to assess the VWFA’s responsivity to words. The localizer run consisted of nine blocks of word presentation interleaved with nine blocks of checkerboard presentation. Each word block lasted 16.5 seconds and contained 15 German words which were presented for 800ms each with an inter-stimulus interval of 300ms. All words consisted of 5-8 graphemes, 2-3 syllables, and were nouns with a word frequency between seven words per million and 10000 words per million as assessed by WordGen (Duyck et al., 2004). During the checkerboard blocks, checkerboard images were presented instead of words. To assess potential rtfMRI NF-driven changes in VWFA responsivity to word stimuli, the functional localizer was repeated a second time after NF training.

### Neurofeedback runs

Each participant underwent a total of six NF training runs. One NF run consisted of four 20-second baseline blocks and four 40-second regulation blocks (see Figure 2). During baseline blocks, participants were instructed to mentally play tennis. This was done to ensure that participants would be engaged in a task unrelated to reading. During the regulation blocks, participants were asked to either upregulate (UP group) or downregulate (DOWN group) their VWFA activity using mental imagery. All participants were informed about the role of the VWFA and its engagement in the reading process. Consequently, participants in the upregulation group performed reading-related mental imagery, and participants in the downregulation group performed mental imagery unrelated to reading. After each run, participants were asked about their exact mental strategy for the respective run. Further, they were asked to estimate their neurofeedback regulation performance and their subjective attention during the run. Finally, participants were given time after each run to think about possible strategies for the next run. This avoided mental strategy planning during baseline periods.

**Figure 2:**
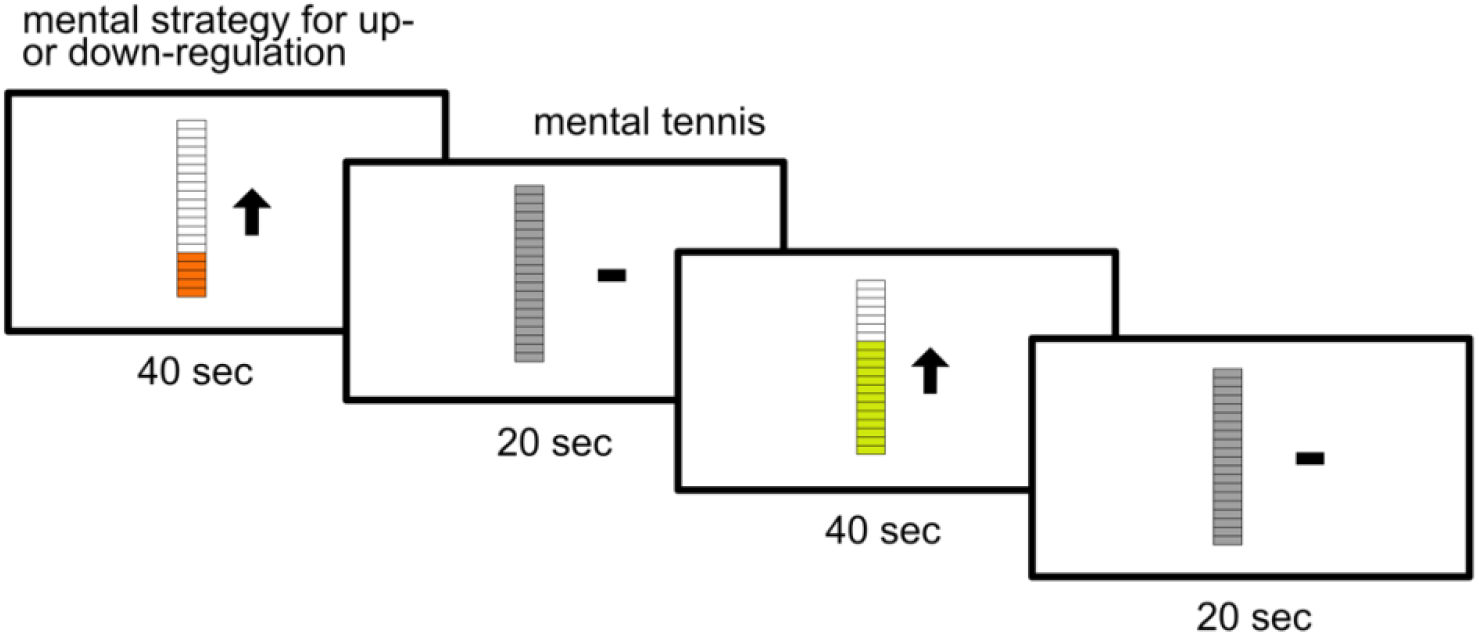
Neurofeedback paradigm. During neurofeedback runs, participants were instructed to either upregulate (UP group) or downregulate (DOWN group) their visual word form area activity using mental strategies. To prevent them from thinking about reading-related strategies during other times, participants were instructed to mentally play tennis during baseline blocks that were interleaved with the regulation blocks.

Participants received continuous visual feedback on their regulation performance in form of a thermometer icon which was filled in based on ongoing VWFA activity levels. In addition, the thermometer icon changed color from red to yellow to green for successful upregulation in the UP group or downregulation in the DOWN group. To emphasize the direction of the regulation task, an upwards (UP group) or downwards (DOWN group) pointing arrow was displayed next to the thermometer icon. During baseline blocks, the arrow was replaced by a neutral line and the thermometer icon turned grey and was static.

### No-feedback runs

Before and after NF runs no-feedback runs were performed. During no-feedback runs, participants had to perform the same mental imagery as during the NF runs, but no feedback was provided. Similarly to the NF paradigm, participants had to perform four 20-second baseline blocks with mental imagery of tennis and four 40-second regulation blocks. Instead of the color-changing thermometer icon, a blue filled thermometer icon was presented during the regulation blocks. Finally, as it was done after each NF run, participants also had to estimate their regulation performance and report their mental strategy and attention after the no-feedback runs. Participants were asked to not perform a new mental strategy in the no-feedback run after NF training, but optimally use the mental strategy which worked best for them during neurofeedback training.

### Image acquisition

The MR session was conducted using a 3 Tesla Philipps Achieva MRI Scanner (Philipps, Best, The Netherlands) located at the MR Center of the Psychiatric University Hospital Zurich. All images were acquired with a 32-channel head coil. The functional VWFA localizer runs each consisted of 247 volumes using a gradient-echo T2*-weighted planar imaging (EPI) sequence with the following parameters: repetition time (TR) = 1250ms, echo time (TE) = 35ms, flip angle (FA) = 80°, field of view (FoV) = 197 × 197 × 138 mm^3^, 42 slices acquired in ascending order, slice gap of 0.3mm, multiband factor 2, and voxel size = 3 × 3 × 3 mm^3^. The first 5 images were discarded as dummy scans. The no-feedback and NF runs were conducted using the same sequence with the following scan parameters: TR = 2000ms, TE = 35ms, FA = 82°, FoV = 220 × 220 × 110 mm^3^, 27 slices in ascending order, 1mm slice gap, no multiband, voxel size of 2 × 2 × 3mm^3^. 130 volumes were acquired per no-feedback/NF run, preceded by an additional five dummy scans which were discarded before data analysis. Finally, a T1-weighted anatomical image was acquired using a FoV of 270 × 255 × 176 mm^3^, voxel size of 1 × 1 × 1 mm^3^, 176 slices, and 9° FA. Extensive descriptions of all scanning parameters can be found in the exam card files uploaded to our Open Science Framework project (https://osf.io/4qhdb/).

### Neurofeedback target ROI

The target ROI for NF training was defined as the 50% most active voxels during a participant’s functional VWFA localizer within a predefined VWFA mask (see Figure S1 in the Supplemental Material). This predefined VWFA mask was created based on ten spheres of varying radii around coordinates taken from the literature (see Supplemental Material Section 1 for a detailed description of the literature coordinates and the creation of the mask) which were chosen to cover a large part of the area associated with the VWFA (The exact mask can be downloaded from our Open Science Framework project: https://osf.io/4qhdb/). For each NF session, the predefined mask was transformed from Montreal Neurological Institute (MNI) space into the participant’s native space. Then, to determine the 50% most activated voxels, the functional VWFA localizer run was analyzed on-site as soon as its image collection was finished. Preprocessing included realignment, coregistration to a single echo planar imaging (EPI) image acquired with the neurofeedback MR sequence right after the localizer run, and smoothing with 6mm full width at half maximum. Normalization was not performed as NF training was performed in native space and because each participant’s target ROI was defined individually. Then, a general linear model (GLM) was calculated with regressors for words, checkerboards, and the six motion parameters. Based on the GLM, a contrast of interest for words versus checkerboards was computed. Finally, this contrast was then thresholded to only receive the 50% most active voxels within the literature-based VWFA mask and subsequently binarized.

### Online analysis of neurofeedback data

Online analysis of the NF runs was performed using the toolbox OpenNFT (Koush et al., 2017) based on MATLAB and Python. OpenNFT received volumes acquired by the MR scanner in real-time using the DRIN export system implemented by Philipps. MR data were preprocessed using OpenNFT’s default preprocessing pipeline including real-time realignment and spatial smoothing. In addition, OpenNFT’s default denoising options were chosen which included the following steps: Drift removal using a cumulative GLM, Kalman filtering for spike removal, and adjustment for serial correlations using a first-order autoregressive model AR(1). Finally, OpenNFT’s default dynamic scaling approach ensured that participants would not reach ceiling or floor effects due to larger changes in signal intensity. After dynamic scaling, the signal was converted to a number between 1 (low brain activity) and 20 (high brain activity) and fed back to the participant in form of the thermometer icon.

### Offline analysis of neurofeedback and no-feedback runs

All offline analyses were performed using SPM12 (Statistical parametric mapping, http://www.fil.ion.ucl.ac.uk/spm/software/spm12/) and MATLAB 2020b (https://www.mathworks.com/products/matlab.html). Offline fMRI data analyses of neurofeedback and no-feedback runs included standard preprocessing steps (slice-time correction, realignment, coregistration, normalization to Montreal Neurological Imaging (MNI) space, and smoothing with 6mm full width at half maximum) for whole brain analyses and preprocessing in native space without normalization and coregistration to the mean functional image instead of the anatomical image for ROI analyses. For both whole brain and ROI analyses, we created a GLM with regressors for regulation and baseline and the six motion regressors. In addition, motion censoring was performed to account for excessive motion (framewise displacement values above 0.9) in single images (Siegel et al., 2014).

For whole brain analyses, the contrast of interest “regulation versus baseline” of the normalized data was used as input data for the second-level GLM. Second-level GLMs were calculated for each group separately across all neurofeedback runs and for a group comparison across all neurofeedback runs. An initial threshold of p = 0.001 was used for single groups and p = 0.005 was used for comparisons between the two groups. Significant clusters were defined based on family-wise error corrections with p < 0.05.

For ROI analyses, the contrast of interest “regulation versus baseline” was defined for the non-normalized data. Then, mean contrast images from the VWFA ROI targeted during NF were extracted for each participant. The mean contrast values were analyzed using mixed model ANOVAs with factors “run number” (pre and post, or neurofeedback run number) and “group” (UP, DOWN), and t-tests where applicable. To account for differences in initial VWFA responsiveness, the mean baseline VWFA activity during the functional localizer was added as a covariate.

### Offline analysis of functional VWFA localizer runs

Functional VWFA localizer runs were preprocessed using realignment, coregistration, normalization to MNI space, and smoothing with 6mm full width at half maximum. An additional preprocessing analysis was performed without normalization to allow for extracting activity levels from the exact ROI that was trained during NF training. For both preprocessed datasets a GLM was calculated using regressors for words, checkerboards, and the six motion parameters. In addition, motion censoring was performed for single volumes with too much motion (framewise displacement values above 0.9). To assess the localization success, whole brain and ROI analyses were performed on a single subject level. The whole brain analysis and ROI analysis were both based on the same “words versus checkerboards” contrast. To assess activity in the right hemisphere, the literature-based mask was flipped from the left to the right hemisphere and contrast values were extracted from this contralateral right hemispheric mask. Paired t-tests were performed to compare the right and left hemispheres. Further, a mixed model ANOVA with factor “run” (PRE, POST) and “group” (UP, DOWN) was used to analyze the extracted mean contrast values of the left mask. Finally, for each group, a second-level analysis with an initial threshold of 0.001 and family-wise error cluster-level correction was performed.

### Statistical analysis of behavioral data

Behavioral data were analyzed using paired and unpaired t-tests, Pearson correlations, and mixed model ANOVAs. Bonferroni correction was performed where applicable. When sphericity was not given for the ANOVAs, corrections were made using Greenhouse-Geisser. All behavioral analyses were performed with IBM SPSS 24 (https://www.ibm.com/de-de/products/spss-statistics) and MATLAB 2020b (https://www.mathworks.com/products/matlab.html).

## Results

### Behavioral and demographic measures before neurofeedback training

The UP and DOWN group did not differ in their IQ scores (t(33)=0.59, p=0.56), working memory (t(38)=1.16, p=0.25), handedness scores (t(37)=0.34, p=0.34), or years of school education (t(37)=1.28, p=0.21). Importantly, the two groups also did not show any significant differences in reading measures as assessed by the SLRT-II and LGVT tests before neurofeedback training. A detailed description of all behavioral measures in the two groups, including several subtests of the two reading tests, is shown in Table 1. Finally, both groups reported similar levels of tiredness (t(38)=1.49, p=0.15), similar scores of motivation to participate in the neurofeedback experiment (t(38)=1.30, p=0.20), and did not differ in their mental imagery or working memory scores.

### Functional localization of the VWFA

The functional VWFA localizer task successfully engaged and localized the VWFA in all participants but one. That one participant demonstrated a negative value when extracting the mean beta value for the “words versus checkerboards” contrast and was, therefore, excluded from further analyses regarding NF. This was necessary because a successful localization of the NF target region in this person could not be achieved. The other participants all showed a highly significant VWFA cluster on a single-subject level (FWE-corrected p<0.001) and positive beta values when extracting the mean beta value for the “words versus checkerboards” contrast from a VWFA mask based on literature (see Methods section and Supplemental Material Section 1 for a description of the literature-based mask). The UP and DOWN group did not differ in their VWFA activity during the localizer task (t(38)=1.00, p=0.32). Further, each participant demonstrated stronger VWFA activity in their left than in their right hemisphere. The difference between left and right VWFA activity was highly significant (t(39)=11.68, p<0.001). An overview of all brain regions engaged during the functional VWFA localizer task before NF in both the UP and DOWN group can be found in the Supplemental Material (Figures S1 and S2, Tables S2 and S3).

### Whole-brain activation during neurofeedback training

On a whole brain level, when contrasting the UP to the DOWN group we found significant (FWE-corrected p<0.05) clusters across the whole reading network including the left precentral gyrus (PCG), the left IFG, the left supramarginal gyrus, the left STG, and the left vOT including the VWFA. In addition, we observed significant activation within the cuneus the supplementary motor area (SMA), the cerebellum, and the orbitofrontal cortex (OFC) (see Figure 3 and Table 2). A detailed description of the results for the UP and DOWN group individually is given in the supplementary material (Tables S4 and S5). One participant of the DOWN group consistently moved their head in one direction, which, over time, resulted in the VWFA moving partly outside the field of view. Therefore, this participant was excluded from further analysis regarding NF.

**Figure 3:**
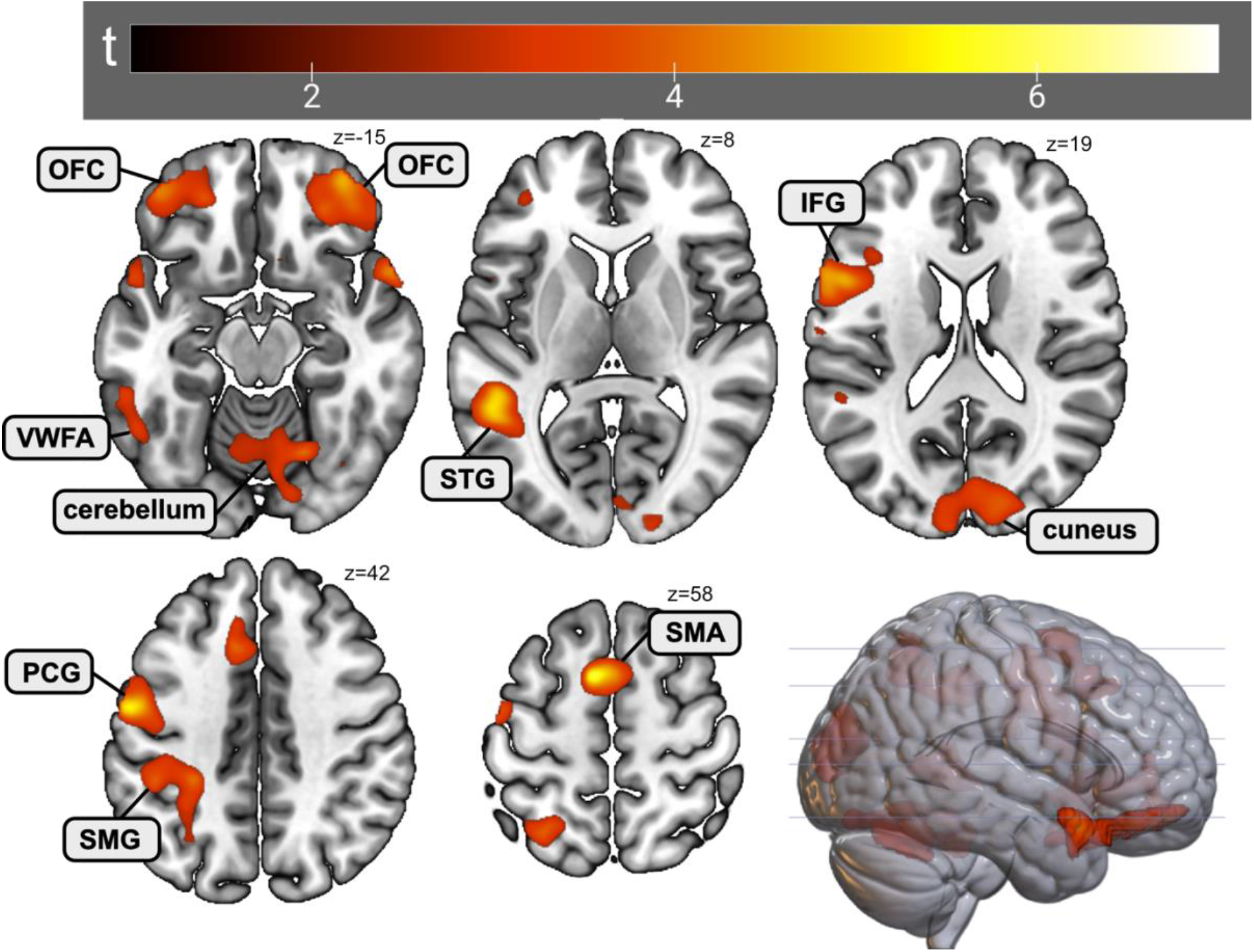
Whole brain activity during neurofeedback runs in the UP group as compared to the DOWN group. When comparing the regulation versus baseline contrast between the two groups, we observed significant (FWE-corrected p<0.05) clusters in the visual word form area (VWFA), the left superior temporal gyrus (STG), the left inferior frontal gyrus (IFG), the left precentral gyrus (PCG), the left supramarginal gyrus (SMG), the cerebellum, orbitofrontal cortex (OFC), cuneus, and the supplementary motor area (SMA).

**Table 2.** Overview of clusters demonstrating higher activity in the UP than in the DOWN group for the regulation versus baseline contrast of neurofeedback runs.

| cluster |  | peak |  | coordinates |  |  | anatomy |
| --- | --- | --- | --- | --- | --- | --- | --- |
| p(FWE-corr) | equivk | p(FWE-corr) | T | x | y | z {mm} |  |
| 0 | 2295 | 0.001 | 6.97 | 24 | -62 | -24 | cerebellum,<br>visual word form area |
|  |  | 0.857 | 3.94 | -18 | -58 | -24 |  |
|  |  | 0.939 | 3.77 | 42 | -62 | -30 |  |
| 0.048 | 726 | 0.007 | 6.25 | -8 | 6 | 60 | supplementary motor area |
|  |  | 0.744 | 4.1 | -12 | 20 | 40 |  |
|  |  | 0.997 | 3.43 | -10 | 30 | 42 |  |
| 0 | 2120 | 0.026 | 5.71 | -56 | -6 | 40 | left precentral gyrus |
|  |  | 0.065 | 5.35 | -48 | -8 | 52 |  |
|  |  | 0.176 | 4.93 | -56 | 2 | 34 |  |
| 0 | 2155 | 0.062 | 5.37 | -42 | 28 | -4 | left inferior frontal gyrus, |
|  |  | 0.367 | 4.57 | -42 | 46 | -12 | left orbitofrontal cortex |
|  |  | 0.386 | 4.55 | -52 | 10 | -6 |  |
| 0.048 | 723 | 0.072 | 5.31 | -52 | -46 | 8 | left superior temporal gyrus |
| 0.001 | 1586 | 0.226 | 4.81 | 36 | 52 | -14 | right orbitofrontal cortex |
|  |  | 0.339 | 4.61 | 54 | 14 | -18 |  |
|  |  | 0.692 | 4.17 | 26 | 18 | -24 |  |
| 0.004 | 1242 | 0.586 | 4.29 | 18 | -86 | 24 | cuneus |
|  |  | 0.833 | 3.98 | -8 | -98 | 16 |  |
|  |  | 0.994 | 3.48 | -8 | -88 | 16 |  |
| 0.01 | 1042 | 0.673 | 4.19 | -28 | -38 | 44 | left supramarginal gyrus |
|  |  | 0.771 | 4.07 | -30 | -60 | 56 |  |
|  |  | 0.826 | 3.99 | -44 | -38 | 40 |  |

### Activation in the Visual Word Form Area during neurofeedback training

For each subject, we extracted mean brain activity within their individual VWFA mask trained during NF. This was done with non-normalized data to replicate activity during online NF training. A mixed model ANOVA with factors group (UP, DOWN) and run (1-6) revealed no significant interaction (F(3.10,108.34)=0.78, p=0.51). Significant main effects were found for the factors group (F(1,35)=5.87, p=0.02) and run (F(3.10,108.34)=3.24, p=0.02), indicating stronger activity in the UP group than the DOWN group and differences in activity between runs (see Figure 4). All ANOVAs were performed using a covariate for VWFA activity during the first functional VWFA localizer run to account for different VWFA baseline activation levels. When excluding this covariate the results remained similar (see Supplemental Material Section 4).

**Figure 4:**
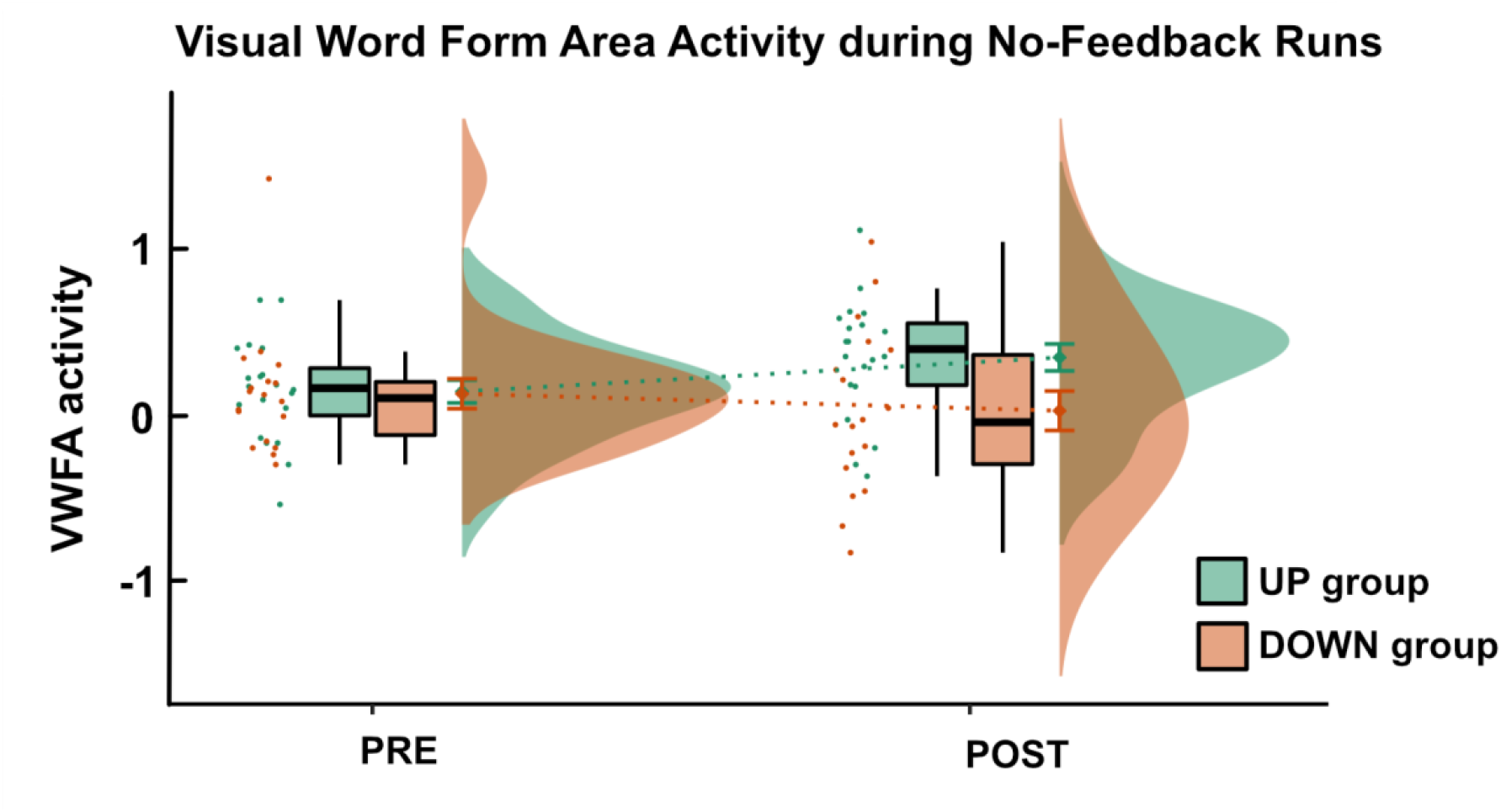
Visual Word Form Area Activity during no-feedback runs before and after neurofeedback training. We observed a significant interaction effect between the factors run (PRE, POST) and group (UP, DOWN). Before neurofeedback training, the two groups did not differ significantly, but after neurofeedback training, the UP group demonstrated higher VWFA activity during the no-feedback run than the DOWN group. Mean values and error bars (depicting one standard error) are displayed on the right side of the boxplots.

### Activation in the Visual Word Form Area during no-feedback runs before and after neurofeedback training

Similarly to the NF analysis, we extracted VWFA activity during no-feedback runs from each subject’s individual NF target region. A mixed model ANOVA with factors group (UP, DOWN) and run (PRE, POST) showed a significant interaction between these two factors (F(1,35)=5.21, p=0.03; Figure 4). Again, baseline VWFA activity during the localizer was used as a covariate (see Supplemental Material Section 4 for results without the covariate). Two-sample t-tests between the two groups showed no significant difference between the groups before NF training (t(36)=0.11, p=0.91), but significantly higher VWFA activity in the UP group as compared to the DOWN group after NF training (t(36)=2.27, p=0.03). When comparing VWFA activity between the PRE and POST time points, we found a significant increase in activity for the UP group (t(19)=2.37, p=0.03), however, no significant change for the DOWN group (t(17)=-0.76, p=0.46)).

### Activation in the Visual Word Form Area during the functional VWFA localizer runs before and after neurofeedback training

Finally, we also extracted VWFA activity during the functional VWFA localizer runs and compared it before and after NF training. Here, we found a significant interaction between the factors group (UP, DOWN) and run (PRE, POST) (F(1,36)=6.81, p=0.01). This interaction showed slightly higher values for the UP than the DOWN group before NF training and lower values for the UP than the DOWN group after NF training (Figure 5). Independent t-tests did, however, not show significant group differences between the UP and DOWN group before (t(36)=0.94, p=0.36) or after (t(36)=-1.66, p=0.11) NF training. In addition, we performed paired t-tests to compare VWFA activity between the PRE and POST time points for each group separately. Here, we found trends for a decrease in VWFA activity for the UP group (t(19)=1.93, p=0.07) and an increase in VWFA activity for the DOWN group (t(17)=-1.79, p=0.09).

**Figure 5:**
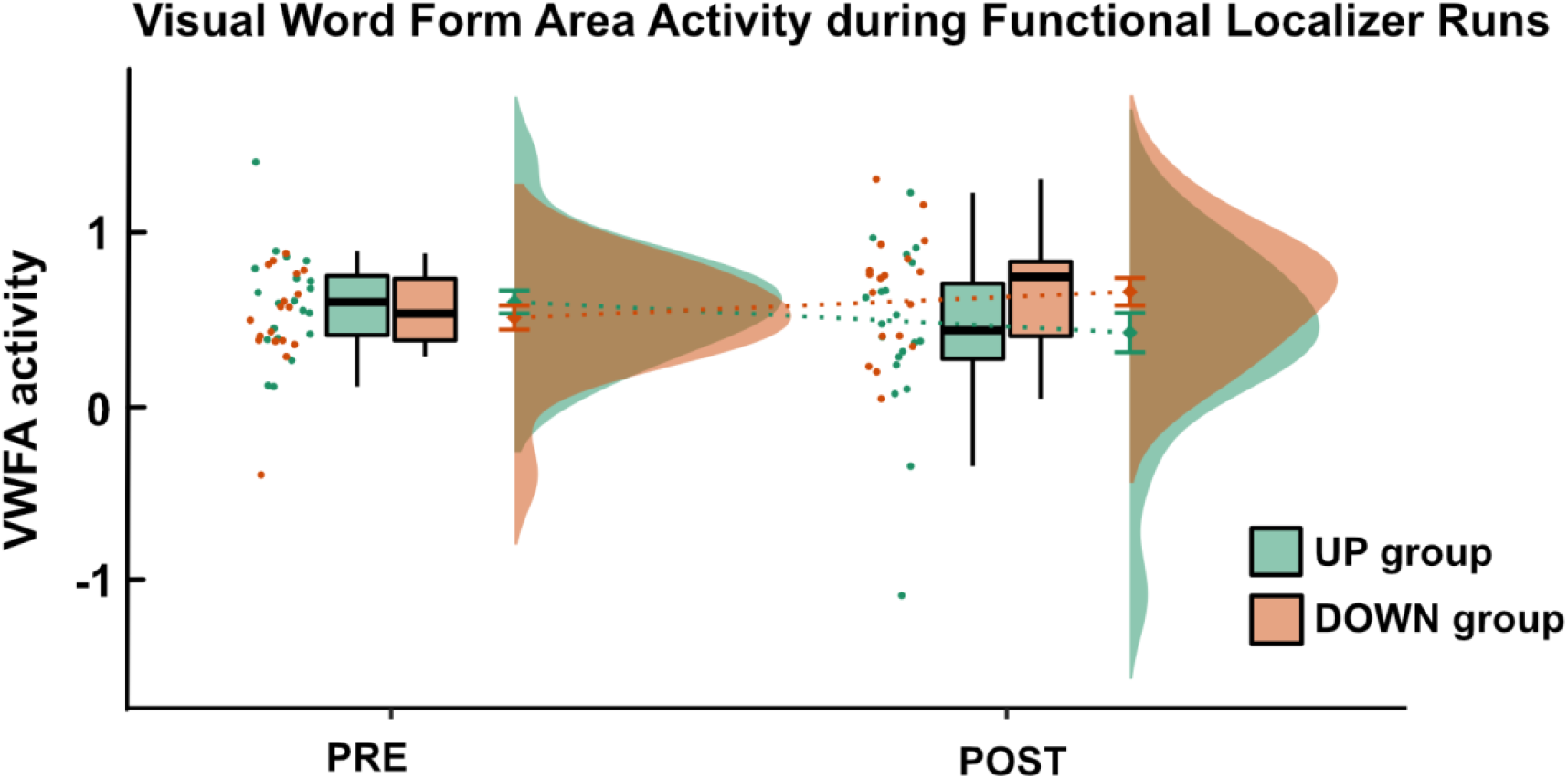
Visual Word Form Area Activity during functional VWFA localizer runs before and after neurofeedback training. We found a significant interaction effect between the factors run (PRE, POST) and group (UP, DOWN) for the functional localizer runs. The two groups did not differ in VWFA activity before or after training, but the UP group demonstrated a trend towards a decrease in activity over time while the DOWN group showed a trend towards an increase in VWFA activity over time.

### Comparison between behavioral reading measures before and after neurofeedback training

For completeness, we also investigated the potential effects of time and/or neurofeedback training on reading skills by performing mixed ANOVAs with factors time (pre, post) and group (UP, DOWN). However, as expected, we did not find any significant interaction effects for reading fluency of words (F(1,34)=0.39, p=0.54) or pseudowords (F(1,34)=0.13, p=0.72), reading comprehension (F(1,38)=0.04, p=0.85), reading speed (F(1,38)=0.17, p=0.68), and reading accuracy (F(1,38)=2.29, p=0.14). However, most measures demonstrated significant improvements over time and we observed a significant main effect of time for reading fluency of words (F(1,34)=22.61, p<0.001) and pseudowords (F(1,34)=20.06, p<0.001), reading comprehension (F(1,38)=18.85, p<0.001), and reading speed (F(1,38)=35.54, p<0.001). Reading accuracy (F(1,38)=0.01, p=0.94) did not show a significant main effect of time. No significant main effect of group was observed for any of the reading measures (see Supplementary Material Section 2 for more details).

## Discussion

Here, for the first time, we investigated the feasibility of regulating brain activity within the VWFA using rtfMRI NF. For the NF runs, our results showed significantly stronger VWFA activity in the participant group who had to upregulate their VWFA as compared to the participant group who received instructions to downregulate their VWFA. Moreover, we did observe significantly higher VWFA activity in the UP than in the DOWN group in the no-feedback transfer run after NF training. These results indicate that self-regulation of the VWFA using mental imagery is indeed feasible.

### Self-regulation of the VWFA

Our NF results clearly demonstrated a significant difference in VWFA activation during NF runs between the UP and DOWN groups. Importantly, the two groups did not show any differences in demographic, cognitive, or reading-related measures. This is in line with previous findings of NF studies observing successful self-regulation in the majority of cases (see (Haugg et al., 2021) for an estimate of NF success in previous studies). However, the two groups differed in their received instructions and corresponding mental imagery.

To investigate the influence of mental imagery performance alone on VWFA activity, we included two no-feedback runs before and after NF training where participants performed the same mental strategies as during NF runs, but no feedback was provided. Importantly, the two groups did not show a significant difference in VWFA activity levels during the no-feedback run prior to NF, but a pronounced difference after NF training. This result nicely shows that, before participants undergo NF training, mental imagery alone seems not sufficient to actively control one’s own VWFA activity.

Consequently, self-regulation of the VWFA can also be performed in the absence of any feedback presentation, but only once this regulation has been practiced, suggesting a successful transfer effect. Importantly, this finding strengthens the potential of VWFA regulation as a therapeutic intervention as it demonstrates that active control of ongoing VWFA activity after NF training and a transfer to everyday life may be possible. This is also in line with other NF studies targeting regions such as the amygdala or the ventral tegmental area that showed successful self-regulation of their respective target areas during no-feedback runs after the NF intervention (MacInnes et al., 2016; Young et al., 2017).

Finally, when comparing VWFA activity during the no-feedback runs before and after NF training for each group independently, we found a significant increase for the UP group, but no change in the DOWN group. Potential floor effects might explain this finding as the strategy of the DOWN group was to think of anything but reading which could already have been successful prior to NF training making it difficult to further downregulate. The UP group, in contrast, most likely had more potential to further improve their mental strategies and to, thus, increase their own VWFA activity.

In conclusion, our findings of the no-feedback runs indicate that self-regulation of one’s own VWFA activity levels is indeed feasible and that, once this self-regulation is learned during NF runs, it can also be performed without the presence of feedback on ongoing VWFA activity levels.

### NF-upregulation vs downregulation training of the VWFA induces activity across the whole reading network

When comparing the UP to the DOWN group for the regulation versus baseline contrast, we did not only find stronger activity in the VWFA, but also in a range of other key regions of the brain’s reading network, including the left STG, left IFG, and left PCG (Martin et al., 2015). This indicates that NF training not only induced the visual processing of mentally imagined orthographic, but most likely also phonological and semantic processing.

In specific, the cluster in the left STG suggests that participants also mentally pronounced the words and letters they visualized mentally. The left STG is associated with both speech perception and speech production (Buchsbaum et al., 2001; Hickok et al., 2000; Yi et al., 2019) and, in the context of reading, has been identified to be involved in the integration of letters and speech sounds (Van Atteveldt et al., 2004). In addition, the left STG has also been found to show hypoactivation in individuals with dyslexia as compared to fluent readers during audiovisual integration tasks, making it another possible target region for NF interventions to improve reading (Blau et al., 2009; Ye et al., 2017). Thus, the additional upregulation of STG activity during NF training might further enhance the potential effects of NF training on reading performance.

Second, we found a significant cluster in the left IFG which is associated with both phonological and semantic processing and has been found to be active during phonological and semantic verbal fluency tasks (Costafreda et al., 2006). This finding also indicates that participants not only visualized the shape of words and letters but also involved phonological and semantic aspects in their mental imagery. When reading, the left IFG is critically involved in semantic integration (J. Huang et al., 2012) and related to phonological awareness and naming ability (P. Turkeltaub et al., 2006). Interestingly, behavioral interventions aimed at improving reading also led to increased activity within the left IFG (Shaywitz & Shaywitz, 2005), again indicating that the additional upregulation of the left IFG during regulation blocks might also enhance potential NF effects of reading skills. Indeed, individuals with dyslexia were found to demonstrate diminished brain activity levels within their left IFG as compared to typical readers (Cao et al., 2006).

Further, we also observed significant activation in the left PCG, a brain region associated with articulatory movements (Baldo et al., 2011) and coordination of speech articulation (Dronkers, 1996). This indicates that some participants may have covertly articulated the imagined letters, words, and sentences during their mental imagery task. As participants were instructed to not move and studies suggest that precentral gyrus activation can be induced by mental imagery alone without explicit overt articulation movements (Mellet et al., 1998), we assume that most participants performed covert articulation rather than explicit articulatory movements.

Finally, we identified a significant cluster in the left SMG. The left SMG, a part of the inferior parietal lobule, has been observed to be involved in visual word recognition (Stoeckel et al., 2009) and phonological processing during reading (Sliwinska et al., 2012). In individuals with dyslexia, the left SMG has been found to show less activation than in typical-reading controls (Ruff et al., 2002). Again, this finding indicates that the enhanced SMG activation during NF training might be an additional factor for improving reading performance even further.

Taken together, our findings indicate that upregulation of the VWFA leads to higher activity in the VWFA, and widespread activity increases across the whole reading network as compared to downregulation. These findings are not surprising as key regions of the reading network have been found to show functional connections related to reading performance (Jasińska et al., 2021; Koyama et al., 2011). Importantly, these activity increases in additional reading-related brain regions might further enhance potential NF-driven improvements in reading performance as many of these other regions, also, have been found to show hypoactivation in individuals with poor reading skills as compared to those with typical reading skills (Martin et al., 2016; Richlan et al., 2009; Yan et al., 2021).

### Further regions demonstrating higher activity levels in the upregulation as compared to the downregulation group during neurofeedback training runs

In addition to regions of the reading network, we also observed other regions with significantly higher activation for the UP than the DOWN group during NF training runs. Some of these regions likely occurred due to the fact that participants of the UP group consistently performed mental strategies related to reading, while mental strategies in the DOWN group differed considerably, making it less likely to consistently engage the same regions in the brain. For instance, some participants in the DOWN group reported having used strategies like “thinking of nothing in specific” or “meditation” while others reported strategies like “reliving memories” or “mental hiking”.

In specific, this difference in mental strategies might explain the significant cluster in regions like the cuneus or supplementary motor area when comparing the UP to the DOWN group. For instance, the cuneus, responsible for visual processing, is more likely to be activated by the visual aspects of reading-related mental strategies than non-visual mental strategies like meditation. In addition, the cuneus has been previously observed to show activation during mental imagery tasks (Stokes et al., 2009). Similarly, it is more likely that the performance of articulatory movements in the UP group consistently induced activity within the supplementary motor area as compared to some of the movement-unrelated strategies of the DOWN group, such as “thinking about nothing”.

### VWFA regulation did not influence reading skills in proficient readers

As expected, we did not find a significant difference in improvements in reading performance between the UP and the DOWN group. In fact, both groups showed significant improvements in their SLRT-II reading fluency scores for words and pseudowords and their LGVT scores for reading comprehension and reading speed. Only reading accuracy did not improve after NF, neither in the UP nor in the DOWN group. The improvement in reading scores in both groups can most probably be explained by practice effects as the same (SLRT-II) or very similar versions of the tests (LGVT) were performed only a few hours after completion of the first tests (Beglinger et al., 2005). The improvement in speed, but not accuracy, indicates that participants tried to be faster in the tests after the NF intervention, a strategy that particularly impacts the performance of the reading fluency test.

Finally, the lack of difference in improvement between the two groups is most likely due to the fact that our participants were adults with typical reading skills who did not demonstrate any reading deficiencies. Indeed, it is rather unlikely that experienced adults without reading impairments would significantly change their reading speed or accuracy solely due to a single session (∼30 mins) of six short instances of VWFA activity upregulation training. Such interventions may be more meaningful in beginning readers or individuals with reading impairments. This is also in line with neuromodulation interventions using transcranial direct current stimulation (tDCS) which showed specific effectiveness in individuals with reading impairments with regards to improvements in reading performance, but not in typical readers (Cancer & Antonietti, 2018; Marchesotti et al., 2020). For instance, Marchesotti and colleagues demonstrated significant improvements in phonological processing and reading accuracy after a tDCS intervention of 20 minutes, but even slightly disturbing effects on individuals without reading impairments (Marchesotti et al., 2020). Consequently, future NF intervention studies targeting the VWFA or other parts of the reading network should either investigate individuals with poor reading skills, beginning readers such as children in the progress of learning to read, or adults learning a new script.

Further, it might also be beneficial to perform NF training over a longer time period to strengthen potential behavioral effects and to benefit from sleep consolidation (Stickgold, 2005). In addition, recent studies have shown that the positive effect of NF training on clinical and behavioral measures increases over time (Rance et al., 2018), most likely due to the fact that, once relevant mental strategies have been learned, participants are able to apply them in their everyday life. As we, also, observed in our study that participants were able to voluntarily increase their VWFA activity during a no-feedback run after NF training, we can assume that these participants would be able to apply their mental strategies over a longer time course outside the MR scanner as well. Consequently, future studies should try to integrate follow-up sessions several days or weeks after the neurofeedback intervention.

### Sustained upregulation of the VWFA led to reduced responsivity to words during passive word viewing

We observed a significant interaction effect between the factors run (PRE, POST) and group (UP, DOWN) for VWFA activity during the localizer runs. Interestingly, this interaction pointed in the opposite direction than expected, namely an increase in VWFA activity from the PRE to the POST run in the DOWN group and a decrease in the UP group. Indeed, paired t-tests demonstrated trends for both a decrease in VWFA activity in the UP group and an increase in VWFA activity in the DOWN group. In comparison, VWFA activity during no-feedback runs increased in the UP group and decreased in the DOWN group. This indicates that NF training enabled participants to actively up- or downregulate their VWFA using mental imagery, but that upregulation would not automatically result in higher VWFA responsiveness during a passive viewing task after the regulation.

One explanation for this finding might be neural adaptation. Previous studies have shown that the VWFA shows neural adaptation to repeated reading of words, i.e. decreased VWFA activity after a repeated reading task (Perrachione et al., 2016; Purcell et al., 2017). Consequently, the consistent activation of the VWFA using reading-related mental imagery in the UP group might have led to decreased responsiveness during passive viewing. Whether or not the participants in the UP group tried to rehearse and mentally read the words that have been presented during the functional VWFA localizer task during their NF regulation blocks is unfortunately not known. Such rehearsal could however explain to some extent the decrease in activation due to repetition and neural suppression effects. In contrast, the DOWN group did not engage their VWFA during mental imagery and, consequently, likely did not influence VWFA responsiveness to orthographic stimuli.

### Limitations

Adults with typical reading skills as participants allowed for investigating the feasibility of self-regulation of the VWFA in this study. Whether or not our results are generalizable to other participant groups and especially to specific patient groups would need to be examined in future studies. In particular, we cannot conclude whether self-regulation of the VWFA is also feasible in people with dyslexia or in children or adults who are in the process of learning to read. Therefore, in future studies, additional participant groups should be included to investigate the feasibility of VWFA regulation in patient populations or during reading development. Further, our adults did not show any deficits in reading performance, making it less likely to achieve considerable improvements in reading performance that are measurable as a result of successful VWFA upregulation.

Second, all reading tasks and MRI tasks were performed within one single session. Consequently, tiredness and attention might have affected task conduction at the end of the session more than at the beginning of the session. Ideally, tasks should be performed before and after NF training at similar fatigue levels. In addition, performing post-training tests on a different day than NF training might allow for sleep consolidation (Stickgold, 2005) which could further improve the effects of NF training. On the other hand, a recent machine learning mega-analysis investigating factors that influence the success of NF studies found no overall beneficial effects of conducting NF studies over several days as compared to just one day (Haugg et al., 2021).

### Outlook

Here, for the first time, we demonstrated the feasibility of self-regulation of the VWFA using rtfMRI NF. As a next step, further studies are needed to investigate whether successful NF training also leads to improvements in reading skills in specific participant groups with poor reading skills or participants who are in the process of learning to read to avoid ceiling effects. As we observed large parts of the reading network to be activated during upregulation as compared to downregulation, the inclusion of additional regions of the reading network might also be possible to boost aspects of the reading process other than visual processing. Finally, future applications of VWFA regulation might also make use of cheaper, more flexible methods such as electroencephalography (EEG), for instance by generating an EEG fingerprint of the fMRI-based VWFA signal and training the fingerprint signal (Keynan et al., 2019; Meir-Hasson et al., 2014).

## Acknowledgments

This study was funded by NCCR Evolving Language, Swiss National Science Foundation, and the Fonds für wissenschaftliche Zwecke im Interesse der Heilung von psychischen Krankheiten.

## Notes

### Competing Interest Statement

The authors have declared no competing interest.

