## Supplemental Material for "Self-Regulation of Visual Word Form Area activation with real-time fMRI neurofeedback"

### Supplementary Material

#### 1. Literature-based Visual Word Form Area Mask

The literature-based VWFA mask was created by defining spheres with different radii around the reported activation peaks of the articles on VWFA listed below using the Marsbar toolbox for SPM (https://marsbar-toolbox.github.io/). These spherical ROIs were then combined to form a joint VWFA mask and used to identify individual ROIs for neurofeedback training and for the analyses of training effects and activation in the functional VWFA localizer task.

| **Literature** | **Coordinates (x,y,z in MNI space)** | **Right hemisphere correlate** | **Radius** |
|---|---|---|---|
| *Impact of literacy on the functional connectivity of vision and language related networks* (López-Barroso et al., 2020) | -44, -50, -14 | 44, -50, -14 | 8 mm |
| *Brain sensitivity to print emerges when children learn letter–speech sound correspondences* (Brem et al., 2010) | -48, −66, −14; | 48, −66, −14; | 6 mm |
| *Variability in Location Impacts Orthographic Selectivity in the "Visual Word Form Area"* (Glezer & Riesenhuber, 2013) | -45, -56, -16 | 45, -56, -16 | 4 mm |
| *Lateralized task shift effects in Broca's and Wernicke's regions and in visual word form area are selective for conceptual content and reflect trial history* (Wallentin, Michaelsen, Rynne, & Nielsen, 2014) | -43, -54, -12 | 43, -54, -12 | 10 mm |
| *The Putative Visual Word Form Area Is Functionally Connected to the Dorsal Attention Network* (Vogel, Miezin, Petersen, & Schlaggar, 2012) | -45, -57, -12 | 45, -57, -12 | 4 mm |

| | | | |
|---|---|---|---|
| *VThe VWFA Is the Home of Orthographic Learning When Houses Are Used as Letters* (Martin et al., 2019) | -34, -55, -13 | 34, -55, -13 | 6 mm |
| *Converging evidence for functional and structural segregation within the left ventral occipitotemporal cortex in reading* (Lerma-Usabiaga, Carreiras, & Paz-Alonso, 2018) | -42, -58, -10 | 42, -58, -10 | 6 mm |

**Table S1. Coordinates and radii of the spheres chosen based on literature.** A general visual word form area mask was created based on previous literature.

*References*

Brem, S., Bach, S., Kucian, K., Guttorm, T. K., Martin, E., Lyytinen, H., … Richardson, U. (2010). Brain sensitivity to print emerges when children learn letter-speech sound correspondences. *Proceedings of the National Academy of Sciences of the United States of America*, *107*(17), 7939–7944. https://doi.org/10.1073/pnas.0904402107

Glezer, L. S., & Riesenhuber, M. (2013). Individual variability in location impacts orthographic selectivity in the "visual word form area." *Journal of Neuroscience*, *33*(27), 11221–11226. https://doi.org/10.1523/JNEUROSCI.5002-12.2013

Lerma-Usabiaga, G., Carreiras, M., & Paz-Alonso, P. M. (2018). Converging evidence for functional and structural segregation within the left ventral occipitotemporal cortex in reading. *Proceedings of the National Academy of Sciences of the United States of America*, *115*(42), E9981–E9990. https://doi.org/10.1073/pnas.1803003115

López-Barroso, D., Thiebaut de Schotten, M., Morais, J., Kolinsky, R., Braga, L. W., Guerreiro-Tauil, A., … Cohen, L. (2020). Impact of literacy on the functional connectivity of vision and language related networks. *NeuroImage*, *213*(March). https://doi.org/10.1016/j.neuroimage.2020.116722

Martin, L., Durisko, C., Moore, M. W., Coutanche, M. N., Chen, D., & Fiez, J. A. (2019). The VWFA is the home of orthographic learning when houses are used as letters. *ENeuro*, *6*(1), 1–13. https://doi.org/10.1523/ENEURO.0425-17.2019

Vogel, A. C., Miezin, F. M., Petersen, S. E., & Schlaggar, B. L. (2012). The putative visual word form area is functionally connected to the dorsal attention network. *Cerebral Cortex*, *22*(3), 537–549. https://doi.org/10.1093/cercor/bhr100

Wallentin, M., Michaelsen, J. L. D., Rynne, I., & Nielsen, R. H. (2014). Lateralized task shift effects in Broca's and Wernicke's regions and in visual word form area are selective for conceptual content and reflect trial history. *NeuroImage*, *101*, 276–288. https://doi.org/10.1016/j.neuroimage.2014.07.012

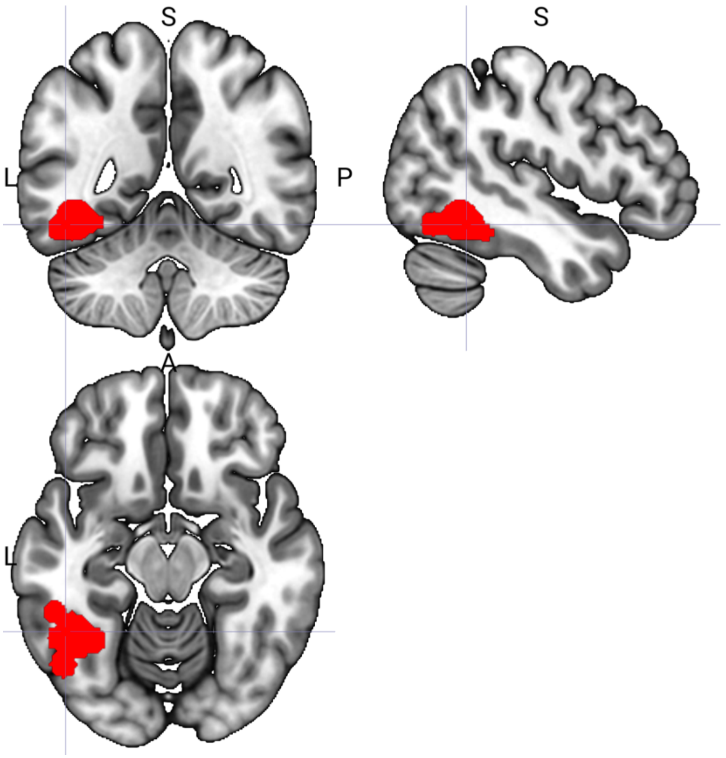

**Figure S1. Literature-based mask of the Visual Word Form Area.** The literature-based Visual Word Form Area (VWFA) mask was created by combining spheres of varying radii with coordinates and radii taken from the literature on the VWFA (Sse table XXXX).

#### 2. Behavioral results

*No main effect of group (UP, DOWN) was found for any of the reading measures*

No significant main effect of group was found for reading fluency of pseudowords ($F(1,34)=2.55$, $p=0.12$ or words ($F(1,34)=1.95$, $p=0.17$), or for reading accuracy ($F(1,38)=0.45$, $p=0.51$), reading speed ($F(1,38)=0.77$, $p=0.39$), and reading comprehension ($F(1,38)=0.25$, $p=0.62$).

#### 3. Functional VWFA localizer results before neurofeedback training (PRE)

*Brain regions engaged by the functional VWFA localizer task in the UP group*

| cluster | | peak | | | | | anatomy |
|---|---|---|---|---|---|---|---|
| p(FWE-corr) | equivk | p(FWE-corr) | T | x | y | z {mm} | |
| 0 | 4022 | 0 | 14.58 | -34 | -92 | -8 | left inferior occipital gyrus, left fusiform gyrus, including VWFA |
| | | 0 | 12.21 | -44 | -66 | -18 | |
| | | 0.004 | 8.51 | -34 | -46 | -20 | |
| 0 | 1904 | 0 | 11.14 | 36 | -86 | -8 | right inferior occipital gyrus |
| | | 0 | 10.69 | 32 | -92 | -2 | |
| | | 0.103 | 6.33 | 4 | -74 | -32 | |
| 0 | 4950 | 0.002 | 8.82 | -42 | 8 | 26 | left precentral gyrus |
| | | 0.019 | 7.43 | -52 | 2 | 28 | |
| | | 0.033 | 7.06 | -50 | 20 | 16 | |
| 0 | 1527 | 0.23 | 5.79 | -48 | -46 | 44 | left supramarginal gyrus, left angular gyrus |
| | | 0.284 | 5.65 | -28 | -58 | 48 | |
| | | 0.385 | 5.42 | -28 | -68 | 56 | |
| 0.004 | 343 | 0.255 | 5.72 | 54 | -10 | 50 | right precentral gyrus |
| | | 0.365 | 5.46 | 38 | -12 | 64 | |
| | | 0.581 | 5.07 | 38 | -28 | 66 | |
| 0.015 | 263 | 0.266 | 5.69 | -28 | -18 | -6 | Left putamen |
| | | 0.292 | 5.63 | -38 | -18 | -20 | |
| | | 0.749 | 4.79 | -28 | -14 | -18 | |
| 0.001 | 487 | 0.396 | 5.4 | -2 | 2 | 72 | supplementary motor cortex |
| | | 0.709 | 4.86 | -8 | -2 | 60 | |
| | | 0.826 | 4.65 | -4 | 26 | 46 | |
| 0.026 | 231 | 0.497 | 5.21 | 50 | -32 | -2 | right middle temporal gyrus |
| | | 1 | 3.6 | 52 | -38 | -10 | |

**Table S2. Whole brain activity during the functional VWFA localizer in the UP group pre training.** Initial threshold $p < 0.001$ uncorrected.

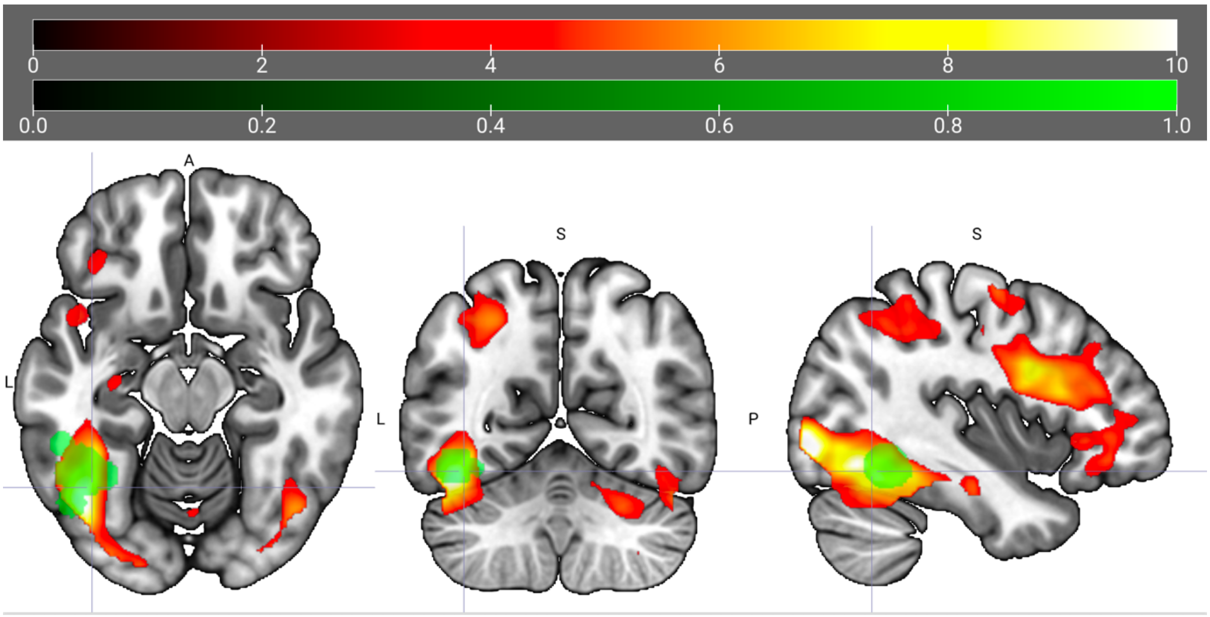

**Figure S1. Whole brain activity during the functional localizer in the UP group pre training.** The Visual Word Form Area mask is highlighted in green. Initial threshold p < 0.001 uncorrected.

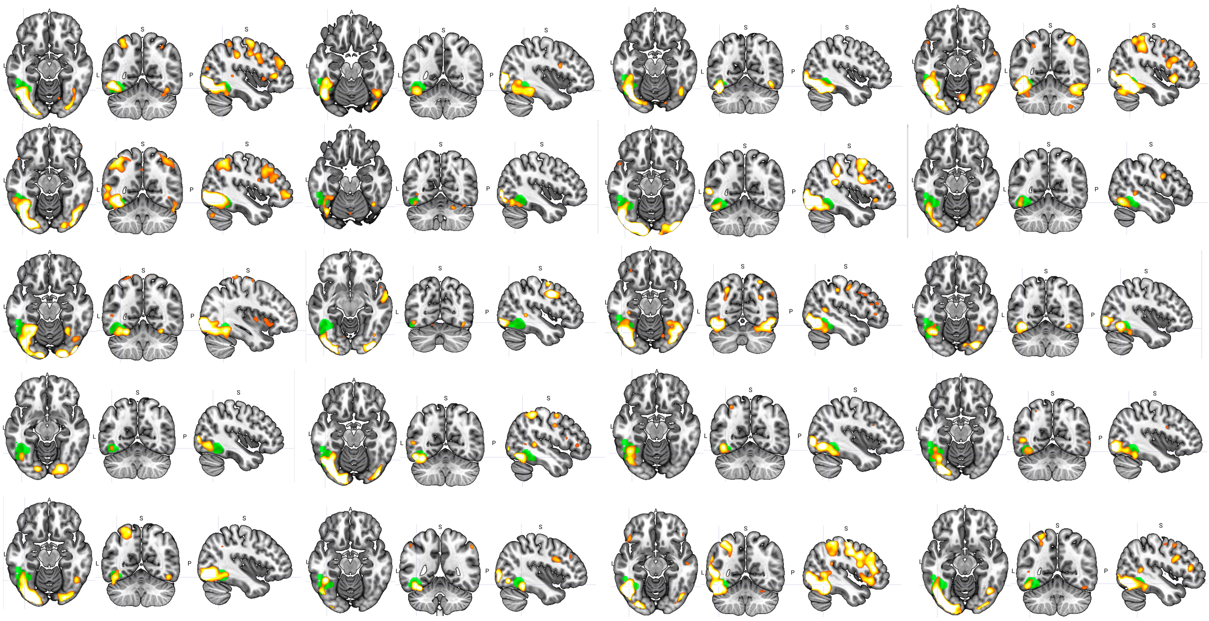

**Figure S2. Individual whole brain activity during the functional VWFA localizer for individuals of the UP group pre training.**

*Brain regions engaged by the functional VWFA localizer task in the DOWN group pre training*

| cluster | | peak | | | | | anatomy |
|---|---|---|---|---|---|---|---|
| p(FWE -corr) | equivk | p(FWE -corr) | T | x | y | z {mm} | |
| 0 | 2696 | 0 | 15.22 | -44 | -64 | -18 | left fusiform gyrus, including VWFA, left inferior occipital gyrus |
| | | 0 | 12.2 | -34 | -88 | -8 | |
| | | 0 | 10.34 | -38 | -46 | -22 | |
| 0 | 707 | 0.002 | 9.39 | -30 | -10 | -14 | left amygdala, left putamen |
| | | 0.635 | 5.11 | -26 | -2 | 10 | |
| | | 0.692 | 5.01 | -38 | -12 | -28 | |
| 0.011 | 270 | 0.011 | 8.08 | 28 | -90 | -4 | right inferior occipital gyrus |
| 0 | 2537 | 0.016 | 7.8 | -36 | 34 | 18 | left middle frontal gyrus |
| | | 0.06 | 6.89 | -44 | 2 | 28 | |
| | | 0.202 | 6.05 | -28 | 4 | 24 | |
| 0 | 635 | 0.354 | 5.63 | -50 | -40 | 38 | left supramarginal gyrus |
| | | 0.532 | 5.29 | -52 | -38 | 46 | |
| | | 0.927 | 4.51 | -44 | -44 | 60 | |
| 0.012 | 262 | 0.426 | 5.48 | 26 | -70 | -44 | right cerebellum |
| | | 0.643 | 5.09 | 18 | -76 | -46 | |
| | | 0.727 | 4.95 | 24 | -70 | -52 | |
| 0.046 | 188 | 0.529 | 5.29 | -8 | 4 | 68 | supplementary motor cortex |
| | | 0.998 | 3.94 | 4 | 2 | 66 | |

**Table S3. Whole brain activity during the functional VWFA localizer in the DOWN group pre training.** Initial threshold $p < 0.001$ uncorrected.

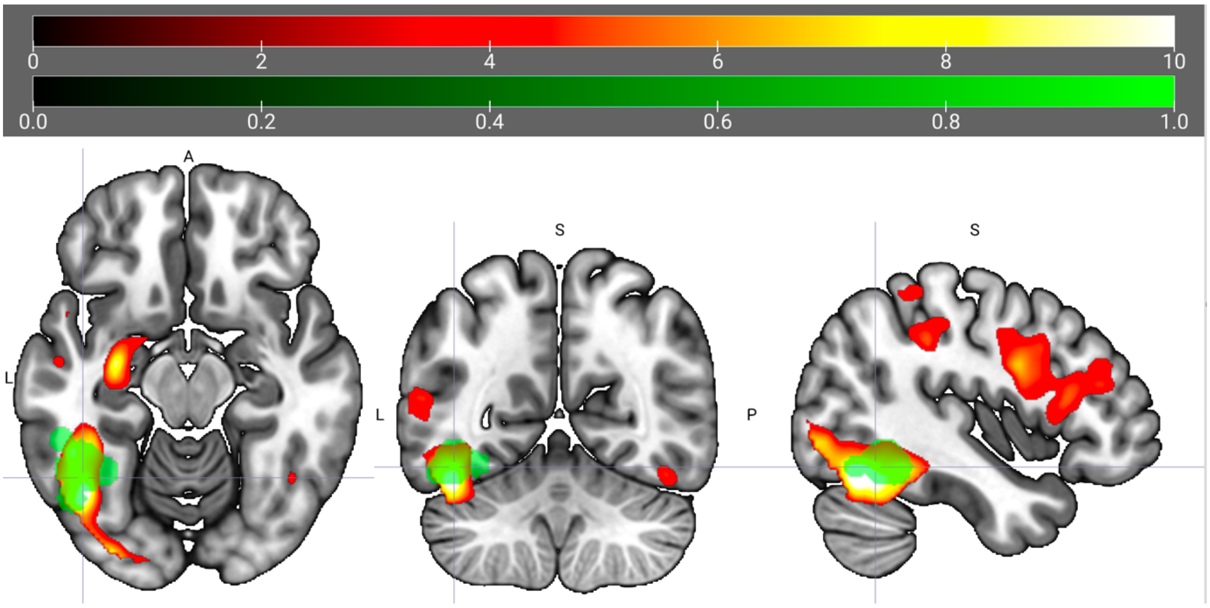

**Figure S3. Whole brain activity during the functional VWFA localizer in the DOWN group pre training.** The Visual Word Form Area mask is highlighted in green. Initial threshold p < 0.001 uncorrected.

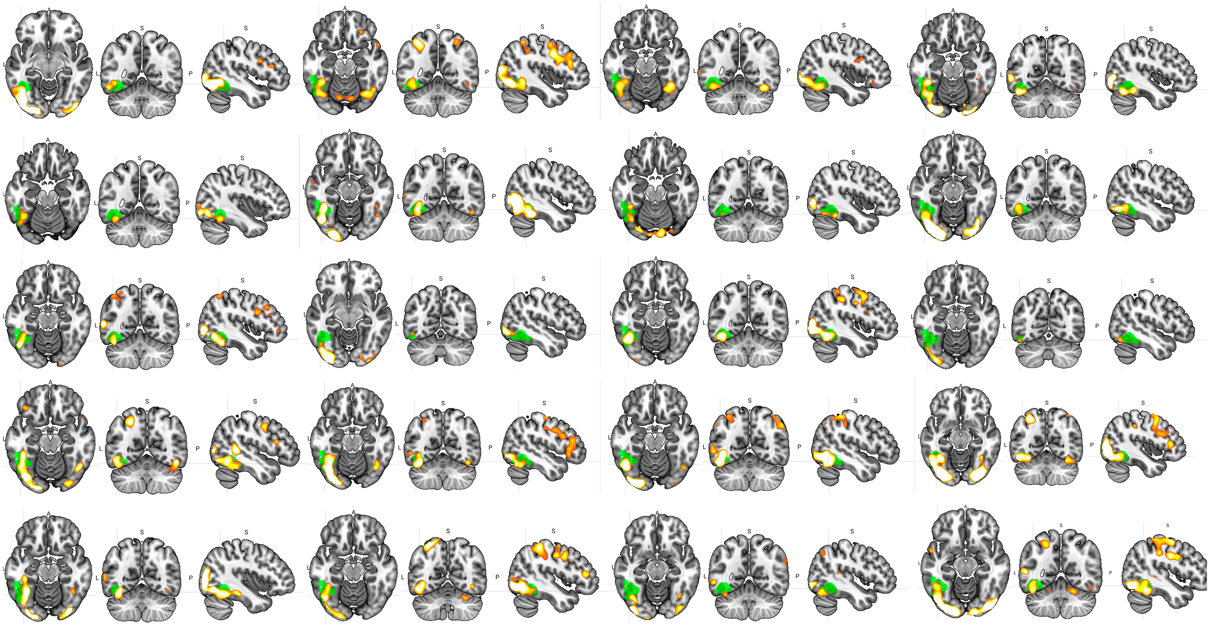

**Figure S2. Individual whole brain activity during the functional VWFA localizer for individuals of the DOWN group pre training.**

*Whole brain group differences between the UP and DOWN group during the functional localizer task prior to neurofeedback training*

We did not observe any significant (cluster corrected) clusters when comparing functional localizer activity between the two groups.

#### 4. Neurofeedback results

*Whole brain activity for the regulation versus baseline contrast in the UP group*

The UP group who was instructed to upregulate their own VWFA activity demonstrated significant clusters across the whole reading network when contrasting regulation to baseline blocks. In specific, we observed significant activity within the bilateral ventral occipital cortex including the VWFA, the bilateral inferior frontal gyrus and bilateral precentral gyrus, the left posterior STG, and the bilateral supramarginal gyrus. In addition, we found significant clusters covering the bilateral anterior insula, the bilateral pallidum, caudate, and ventral tegmental area, and the bilateral supplementary motor area and superior parietal lobule (see Table S4 for detailed coordinates).

| **cluster** | | **peak** | | **coordinates** | | | **anatomy** |
|---|---|---|---|---|---|---|---|
| p(FWE-corr) | equivk | p(FWE-corr) | T | x | y | z {mm} | |
| 0 | 9030 | 0 | 12.65 | 34 | 26 | -2 | right anterior insula<br>right IFG<br>including right precentral gyrus |
| | | 0 | 10.39 | 50 | 12 | 18 | |
| | | 0 | 10.16 | 46 | 14 | 26 | |
| 0 | 7111 | 0 | 10.08 | -40 | 24 | -4 | left IFG<br>including left precentral gyrus<br>left anterior insula |
| | | 0 | 10.07 | -38 | 32 | -2 | |
| | | 0.001 | 9.37 | -30 | 24 | -6 | |
| 0 | 6506 | 0.001 | 9.3 | 40 | -72 | 18 | occipital gyrus<br>right fusiform gyrus |
| | | 0.004 | 8.3 | 46 | -66 | 10 | |
| | | 0.006 | 8.08 | 44 | -64 | -10 | |
| 0 | 1889 | 0.011 | 7.69 | -8 | 20 | 44 | left supplementary motor area |
| | | 0.014 | 7.56 | -8 | 8 | 58 | |
| | | 0.014 | 7.52 | 6 | 20 | 42 | |
| 0 | 1195 | 0.012 | 7.63 | -24 | -54 | 44 | left superior parietal lobule, left supramarginal gyrus |
| | | 0.107 | 6.22 | -24 | -66 | 48 | |
| | | 0.376 | 5.35 | -44 | -40 | 40 | |
| 0.003 | 487 | 0.06 | 6.59 | -44 | -66 | -8 | left ventro-occipital cortex (VWFA) |
| | | 0.488 | 5.14 | -46 | -56 | -12 | |

| | | | | | | | |
|---|---|---|---|---|---|---|---|
| 0.009 | 387 | 0.163 | 5.94 | -52 | -46 | 8 | left posterior STG |

**Table S4: Whole brain activity for the regulation versus baseline contrast in the UP group**

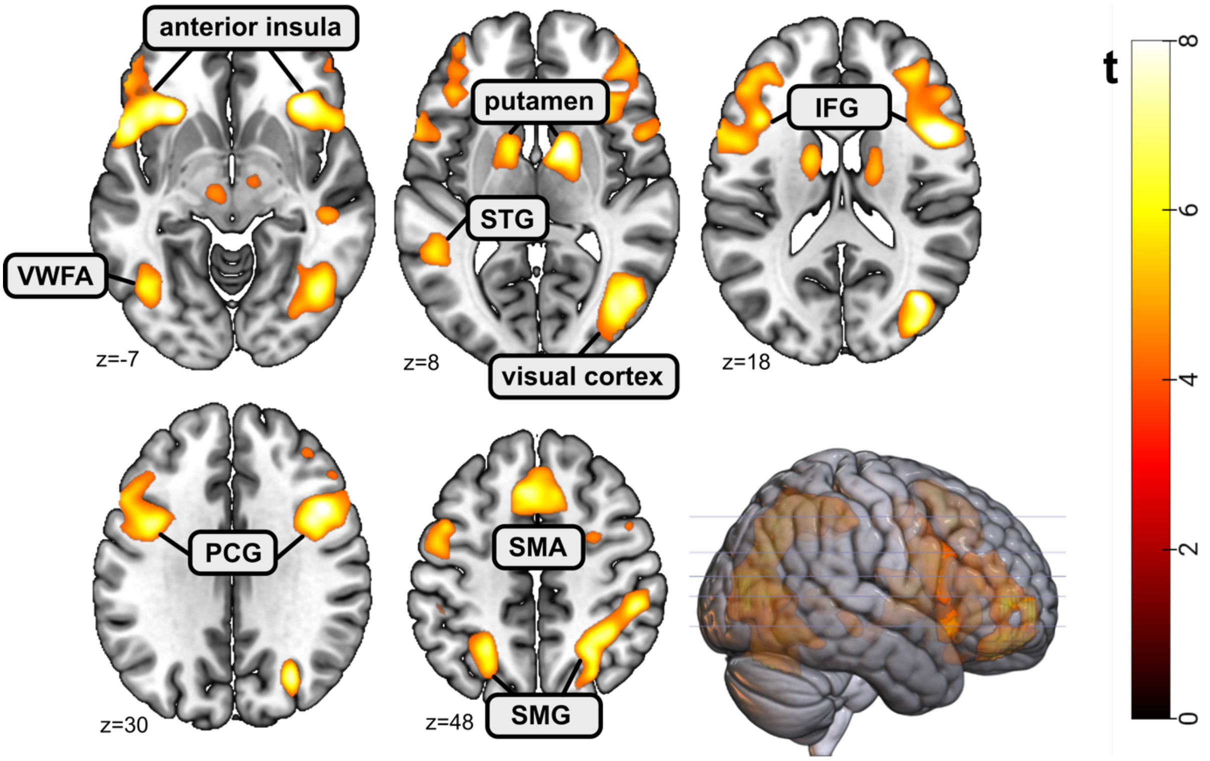

**Figure S5: Whole brain activity for the regulation versus baseline contrast in the UP group.** Initial threshold p < 0.001 uncorrected. Abbreviations: visual word form area (VWFA), superior temporal gyrus (STG), inferior frontal gyrus (IFG), precentral gyrus (PCG), supplementary motor area (SMA), supramarginal gyrus (SMG).

*Whole brain activity for the regulation versus baseline contrast in the DOWN group*

In the DOWN group who was instructed to downregulate their VWFA activity we observed significant activity in the right ventral occipital cortex, the right superior parietal lobule, the right supramarginal gyrus, the right precentral gyrus, the right inferior frontal gyrus, the right supplementary motor area, and the bilateral anterior insula (see Table S5 for detailed coordinates).

| cluster | | peak | | | | | | |
|---|---|---|---|---|---|---|---|---|
| p(FWE -corr) | equivk | p(FWE -corr) | p(FDR -corr) | T | x | y | z {mm } | anatomy |
| 0 | 4164 | 0 | 0.002 | 11.28 | 46 | 10 | 22 | right precentral gyrus right inferior frontal gyrus |
| | | 0.009 | 0.013 | 8.34 | 38 | 38 | 28 | |
| | | 0.018 | 0.017 | 7.81 | 30 | 24 | 2 | |
| 0 | 4961 | 0.001 | 0.004 | 10.13 | 46 | -68 | -10 | right inferior occipital gyrus right superior parietal lobule right supramarginal gyrus |
| | | 0.004 | 0.009 | 8.94 | 26 | -60 | 46 | |
| | | 0.005 | 0.009 | 8.85 | 38 | -86 | 18 | |
| 0.018 | 342 | 0.129 | 0.079 | 6.37 | 6 | 14 | 54 | right supplementary motor area |
| | | 0.335 | 0.151 | 5.64 | 8 | 28 | 42 | |
| 0.046 | 266 | 0.372 | 0.151 | 5.55 | -28 | 22 | 0 | left anterior insula |

**Table S5: Whole brain activity for the regulation versus baseline contrast in the DOWN group**

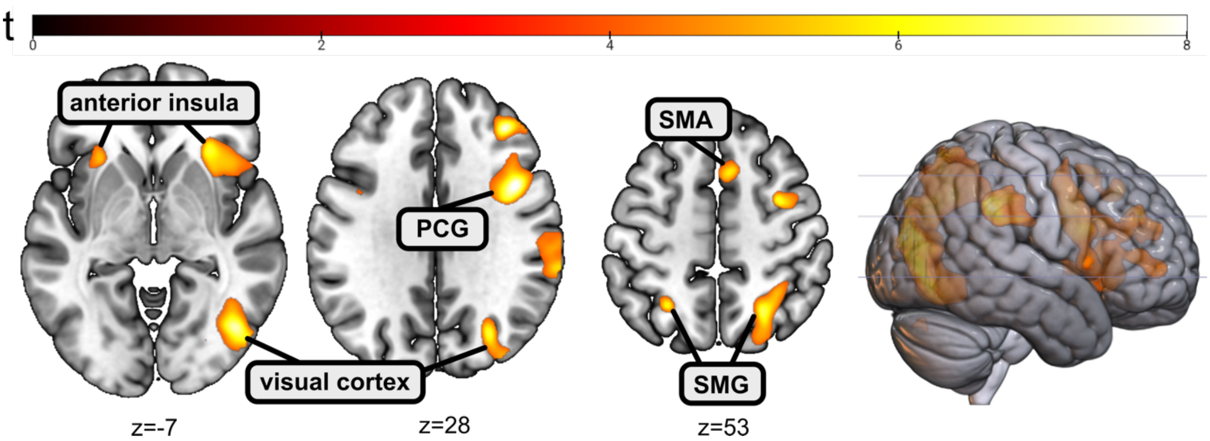

**Figure S6: Whole brain activity for the regulation versus baseline contrast in the DOWN group.** Initial threshold p < 0.001 uncorrected. Abbreviations: precentral gyrus (PCG), supplementary motor area (SMA), supramarginal gyrus (SMG).

*Activation in the Visual Word Form Area during neurofeedback training without a correction for baseline Visual Word Form Area activity levels during the localizer run*

A mixed model ANOVA with conditions group (UP, DOWN) and run (1-6) revealed no significant interaction ($F(3.16,113.74)=0.90$, $p=0.45$). A significant main effect was found for both the condition run ($F(3.16,113.74)=0.05$, $p<0.001$), and the condition group ($F(1,36)=5.13$, $p=0.03$).

*Activation in the Visual Word Form Area during no-feedback runs without a correction for baseline Visual Word Form Area activity levels during the localizer run*

A mixed model ANOVA with conditions group (UP, DOWN) and run (1-6) revealed a trend for an interaction ($F(1,36)=3.77$, $p=0.06$). No significant main effect was found for both the condition run ($F(1,36)=0.40$, $p=0.53$) and the condition group ($F(1,36)=2.79$, $p=0.10$).
